# *MaiZaic*: a robust end-to-end pipeline for mosaicking freely flown aerial video of agricultural fields

**DOI:** 10.1101/2024.12.31.630534

**Authors:** Dewi Endah Kharismawati, Toni Kazic

**Affiliations:** Dept. of Electrical Engineering and Computer Science, University of Missouri, Columbia, MO; Missouri Maize Center, University of Missouri, Columbia, MO; Interdisciplinary Plant Group, University of Missouri, Columbia, MO; Institute for Data Science and Informatics, University of Missouri, Columbia, MO

## Abstract

Unmanned aerial vehicles (UAVs) are increasingly used for high throughput phenotyping. In principle, freely flown vehicles would permit real-time flexibility in identifying and scouting regions of interest. Mosaicking multiple images provides a high resolution global image and consumer-grade UAVs offer low cost, ease of flying, and excellent RGB cameras. The vehicles’ inaccurate telemetry complicates estimating the homographies between pairs of frames, the standard mosaicking approach. Moreover, errors accumulate during computation, distorting later portions of the mosaic. Finally, crop fields are particularly challenging to mosaic because their planting is so regular and the plants are so similar, eliminating distinctive features that could guide mosaicking. We propose *MaiZaic*, an end-to-end pipeline that dynamically samples video frames using optical flow, automates camera and gimbal calibration, estimates homographies with an unsupervised convolutional neural network, detects shots among frames, and generates mini-mosaics. Together, these techniques significantly reduce errors in the output mosaics. Our deep learning model is trained on a comprehensive video dataset comprising different flight trajectories, maize lines, growth stages, and augmented illumination data. *MaiZaic* is more accurate and faster than ASIFT and more robust than our earlier *CorNet* and *CorNetv2*. We demonstrate *MaiZaic*’s effectiveness in generating accurate mosaics of imagery captured by freely-flown UAVs and explore its generalizability.

**Core ideas:**

- *MaiZaic* is an end-to-end pipeline to mosaic freely flown agricultural imagery captured with consumer-grade UAVs.
- *MaiZaic* introduces novel algorithms that efficiently choose video frames, calibrate, and mosaic the imagery.
- Our unsupervised deep homography estimator, *CorNetv3*, is 14 times faster and 8.59% more accurare than ASIFT.
- *MaiZaic* generalizes well and mosaicks maize at different growth stages, objects, trajectories, cameras, and pilots.
- The mini-mosaicking algorithm improves mosaic accuracy by interrupting error accumulation.

## 1 INTRODUCTION

The recent explosion in phenotyping and precision agriculture offers new opportunities for monitoring plants, improving decision making, and automating management during field seasons. Rather than scouting by walking through fields and inspecting plants in the customary, labor-intensive way, Unmanned Aerial Vehicles (UAVs) offer a very accessible method to obtain high resolution data. Consumer-grade vehicles offer low cost, easy flying, and remarkably superb RGB cameras in an effective package (Araus & Cairns, 2014; Minervini, Scharr, & Tsaftaris, 2015; Feng, Zhou, Vories, Sudduth, & Zhang, 2020; Aktar et al., 2020; Kharismawati et al., 2020; Vong, Conway, Zhou, Kitchen, & Sudduth, 2021; Zheng et al., 2021; Kharismawati et al., 2021; Kharismawati, Bunyak, Palaniappan, & Kazic, 2022). Two types of trajectories can be flown: capturing still images while hovering over a set of predetermined waypoints at fixed altitude; and recording video while freely flying a trajectory that can vary in altitude, path, and speed. The freely flown trajectory can be immediately adapted in flight to meet the challenges a researcher’s or farmer’s crop faces.

Flying at low altitudes captures a field as high resolution patches that must be mosaicked together for a comprehensive view of the whole field. However, mosaicking agricultural imagery is very challenging. Nearly all row crops and orchards are planted in regular patterns as monocultures, so that to the computer all the plants look nearly alike and there is little to distinguish different parts of a field. Mosaicking relies on somehow finding regions in two images that are very similar. These similarities are expressed as *feature descriptors*, multidimensional vectors that capture image textures as places where gradients of color, entropy, or other properties intersect or rapidly change (Bay, Tuytelaars, & Van Gool, 2006; Morel & Yu, 2009; Juan & Gwun, 2009; Aktar, Aliakbarpour, Bunyak, Seetharaman, & Palaniappan, 2018; Aktar et al., 2020). Constructing, evaluating, and matching feature descriptors is computationally expensive. To let Structure from Motion and bundle adjustment algorithms compute mosaics in reasonable time, commercial software such as Pix4D, Agisoft, and WebODM simplify the mosaicking problem by requiring a fixed flight trajectory that guarantees extensive image overlap (Pix4DMapper, 2017; Schonberger & Frahm, 2016; Yanagi & Chikatsu, 2016; Agisoft LLC, 2019; OpenDroneMap, 2020; Kameyama & Sugiura, 2021; Geospatial Technology and Applications Center, 2022; Mora-Félix, Rangel-Peraza, Monjardín-Armenta, & Sanhouse-García, 2024). These packages also benefit from installing ground control points in the field, an especially labor-intensive task for repeated imaging throughout the season (Ferrer-González, Agüera-Vega, Carvajal-Ramírez, & Martínez-Carricondo, 2020). When not enough features are matched, for example because of poor telemetry or variable image overlap, these approaches struggle and erroneously interpolate from the neighboring features, producing artifactual mosaics with “melted” areas or missing pixels (Supplemental Figure 9).

Another approach to mosaicking is to compute *homographies* — a set of affine transformation matrices that map feature descriptors from one image to their corresponding features in another, even when the two images are taken from different perspectives (Fischler & Bolles, 1987; Aktar et al., 2016; DeTone, Malisiewicz, & Rabinovich, 2016; Schönberger & Frahm, 2016; Ufuktepe et al., 2020; Barath, Mishkin, Polic, Förstner, & Matas, 2023). To build a mosaic, homography matrices are computed between successive pairs of images, using the matrix for each pair to transform the second image into the reference frame of the first. A transformed image is laid into the growing mosaic by computing the homography between that image and the product of the previous homography matrices:

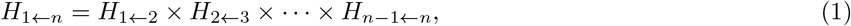

where *H* is an homography matrix, the subscript *i* ← *j* denotes the homography matrix for frame *j* onto frame *i*, and *H*_1←*n*_ is the accumulated homography matrix to project frame *n* onto frame 1. Since the homography matrices account for changes in lateral motion (translation), rotation, and scaling (altitude changes) without using metadata or extrinsic camera poses, they can mosaic imagery from freely flown, consumer-grade UAVs. However, estimating homography is a slow and computationally expensive process that scales up with image number and resolution. Moreover, tiny errors in the homographies accumulate during mosaicking from the product term, so that frames of the same spot viewed at two different times during the flight can be misaligned.

We previously addressed the problems of inflexible trajectories and slow computation by developing *CorNet*, an unsupervised convolutional neural network that estimates homographies from video imagery captured by freely flown UAVs without using metadata or ground control points (Kharismawati et al., 2020). *CorNet* estimates the homography matrices more accurately and ten times faster than ASIFT, the gold standard feature descriptor (Morel & Yu, 2009). However, this early version of *CorNet* struggles to mosaic fields of seedlings; inappropriately reuses camera errors; accumulates errors during mosaicking; and performs poorly when the field’s illumination changes among the frames, for example when clouds move over the field, shadowing parts of it.

To address these problems while producing more accurate mosaics even faster, we introduce *MaiZaic*, a pipeline that incorporates five significant improvements. First, we minimize the number of frames needed for the mosaic by developing a dynamic sampling algorithm to select frames that have the optimal overlap, choosing fewer frames from the straight parts of the trajectory and more when the UAV turns. Second, we automatically calibrate the lens and gimbal to correct errors occurring during flight, improving the quality and accuracy of the global mosaic. Third, we retrain the *CorNet* homography estimation model using an expanded dataset that includes serpentine trajectories, multiple plant growth stages, and synthetic ilumination changes. Fourth, we implement an improved shot detection algorithm optimized for complex trajectories to divide frames into shots, since the classic graphcut algorithm for shot detection performs well only for simple, orbital trajectories (Viguier et al., 2015; Farnebäck, 2003). Fifth, we reduce error accumulation along the mosaic by computing mini-mosaics for each shot (Aktar, Aliakbarpour, Bunyak, Kazic, et al., 2018; Aktar et al., 2020). A companion paper describes the computational details of these improvements (Kharismawati & Kazic, 2025). These improvements have been assembled into an end-to-end pipeline that takes an aerial video and produces a global mosaic from freely flown UAV trajectories. We have explored *MaiZaic*’s generalizability over different objects, trajectories, cameras, vehicles, and pilots. Together, these improvements bring automation of efficient dynamic scouting by inexpensive UAVs closer to reality.

## 2 MATERIALS AND METHODS

### 2.1 Materials

#### Maize and Other Fields

Three inbred, an elite, and multiple disease lesion mimic mutant lines of *Zea mays* were planted in genetic nurseries in Columbia, Missouri, USA in the 2021, 2022, and 2023 summer field seasons. The field layout was the same in all years: the 6.1 m long rows were planted 0.9 m apart with plants spaced ≈12–24.5 cm apart depending on the line. The ranges, sets of parallel rows that span the width of the field, were separated by unplanted 1.2 m alleys for easier access. The center file of rows was left unplanted, except for one row in 2023. All nursery rows were hand planted and the surrounding border of elite maize was machine planted. The production fields were located in Sumberarum, East Java, Indonesia in winter 2024. There maize is typically planted in rows with a spacing of 70 cm between rows and 20–25 cm between plants within each row. All fields were hand-planted. We also flew fields of pumpkins and sunflowers and groups of farm buildings in the 2023 field season.

#### Flights

A 2021 DJI Mavic 2 Pro equipped with an RGB camera and a 2.54 cm CMOS sensor was manually flown along several trajectories with either nadir (perpendicular to the earth’s surface) or an oblique camera pose. In all trajectories the UAV was flown at varying altitudes and speeds. The first trajectory was a serpentine perpendicular to the rows, translating the UAV without rotation between passes from one range to the next (a *slide*), so that relative to the direction of motion, the camera looks backward in one pass and forward in the next. Each pass captures a range at an average speed of 1.5–3.5 kph and an altitude of 7.5–15 m above the ground. The second trajectory was a serpentine parallel to the rows with slides between passes. The altitude varied, depending on the number of passes and the width of the field, and usually ranged from 12–25 m above the ground. The third trajectory involved rotation (clockwise and counterclockwise), scaling (altitude changes), and sideways motions (left and right) around the Indonesian maize field. For the pumpkin and sunflower fields, we flew a serpentine parallel to the edge of the field. The last trajectory is an unconstrained cruise around the Missouri nursery field and farm buildings. To explore *MaiZaic* generalizibility, we included test data collected with our secondary UAV, a DJI Air 2S operated by a newly trained pilot, and the UMCD dataset, capured using a DJI Phantom 3 (Avola et al., 2020).

#### Video Data and Model Training

Video from each Mavic 2 Pro flight was captured at a resolution of 3840 × 2160 pixels, while the Air 2S recorded with a resolution of 5472 × 3078 pixels, both saving their videos in .MOV format at a frame rate of 30 fps. The lenses of the UAVs were calibrated using a 1400 × 1400 × 6 mm checkerboard with 12 × 9 squares purchased from calib.io, Odense, Denmark. This was fastened to a 1.90 cm thick piece of plywood to increase its planarity.

The *CorNetv2* deep learning model was trained on one NVIDIA GeForce GTX 1080 Ti GPU processor with 11GB memory in the university’s cluster. The *CorNetv3* deep learning model was trained on a Lambda Labs machine using its two NVIDIA RTX 2080Ti GPUs each with 11GB; this machine also has an Intel Core i9-9920X CPU and 128 GB of RAM. The whole pipeline was deployed on this machine. Table 1 summarizes the training, and Table 2, the test videos used in this paper.

**Table 1:** Training data extracted by dynamic sampling from imagery captured by our DJI Mavic 2 Pro. The default sampling rate was 2 fps for all.

| crop/date/video_id | trajectory (no. passes) | pose | growth stage | no. frames/video |
| --- | --- | --- | --- | --- |
| 21r/23.7/412 | backward (1) | nadir | seedlings | 191 |
| 21r/27.7/427–429 | parallel (4) | nadir | seedlings | 620,624,261 |
| 21r/6.8/451 | parallel | nadir | adult | 60 |
| 21r/14.8/480 | single plant orbit | oblique | adult | 337 |
| 21r/16.8/527–528 | perpendicular | nadir | adult | 636,587 |
| 21r/19.8/613 | parallel (1) | nadir | adult | 364 |
| 21r/19.8/614 | parallel, forward (1) | nadir | adult | 389 |
| 21r/1.9/692 | parallel, forward (1) | nadir | adult | 386 |
| 21r/1.9/693 | parallel, backward (1) | nadir | adult | 562 |
| 21r/1.9/694–696 | perpendicular | oblique | adult | 630,635,25 |
| 21r/15.9/716 | backward | nadir | adult | 240 |
| 21r/15.9/717–718 | perpendicular | nadir | adult | 589,469 |
| 22r/16.6/894 | forward (1) | nadir | seedlings | 208 |
| 22r/16.6/896 | satellite view, still | nadir | seedlings | 55 |
| 22r/16.6/917 | perpendicular | nadir | seedlings | 677 |
| 22r/16.6/923–924 | forward (1) | nadir | seedlings | 169,150 |
| 22r/17.6/925–928 | perpendicular, rotation | nadir | seedlings | 777,832,802,473 |
| 22r/17.6/933–934 | parallel (4) | nadir | seedlings | 702,286 |
| 22r/18.6/941–943 | parallel, rotation (many) | nadir | seedlings | 776,741,223 |
| 22r/18.6/944 | high altitude parallel (1) | nadir | seedlings | 258 |
| 22r/18.6/945 | parallel, rotation (2) | nadir | seedlings | 607 |
| 22r/18.6/956 | parallel, rotation (2) | nadir | seedlings | 548 |
| 22r/21.6/1–2 | perpendicular | oblique | seedlings | 1179,161 |
| 22r/22.6/996–999 | perpendicular, slide | nadir | seedlings | 703,666,706,15 |
| 22r/22.6/1 | perpendicular, higher | nadir | seedlings | 487 |
| 22r/24.6/38 | perpendicular | oblique | seedlings | 449 |
| 22r/14.7/228 | forward (1) | nadir | adult | 154 |
| 22r/14.7/229–235 | perpendicular | nadir | adult | 779,169,79,98,620,663,251 |
| 22r/19.7/283 | forward | nadir | adult | 223 |
| 22r/19.7/284 | high altitude | nadir | adult | 30 |
| 22r/5.8 /393–396 | perpendicular, slide | nadir | adult | 739,646,632,509 |
| 22r/5.8 /397 | forward | nadir | adult | 128 |
| 22r/10.8/508 | high altitude | nadir | adult | 31 |
| 22r/19.8/669 | parallel (1) | nadir | adult | 177 |
| 24n/17.12/221–224 | rotation | nadir | seedling | 1107,1190,1228,961 |
| 24n/17.12/225–228 | sideways | nadir | seedling | 792,619,583,521 |
| 24n/17.12/232–235 | scale | nadir | seedling | 547,887,827,180 |
| 24n/18.12/237–239 | rotation | nadir | adult | 584,772,267 |
| 24n/18.12/241–242 | sideways | nadir | adult | 754,285 |
| 24n/20.12/243–246 | rotation | nadir | adult | 1207,1153,1087,660 |
| 24n/20.12/247 | sideways | nadir | adult | 838 |
| 24n/20.12/248–250 | sideways | nadir | seedling | 564,668,608 |
| 24n/20.12/252 | scale | nadir | adult | 1055 |
| 24n/20.12/253–255 | scale | nadir | seedling | 946,1046,886 |
| 24n/21.12/315–316 | rotation | nadir | adult | 1256,976 |
| 24n/21.12/317–318 | rotation | nadir | seedling | 1214,690 |
| 24n/21.12/319–323 | sideways | nadir | adult | 856,916,52,772,784 |
| 24n/21.12/325 | scale | nadir | adult | 1083 |
| 24n/21.12/326–328 | scale | nadir | seedling | 1096,1095,202 |

**Table 2:**
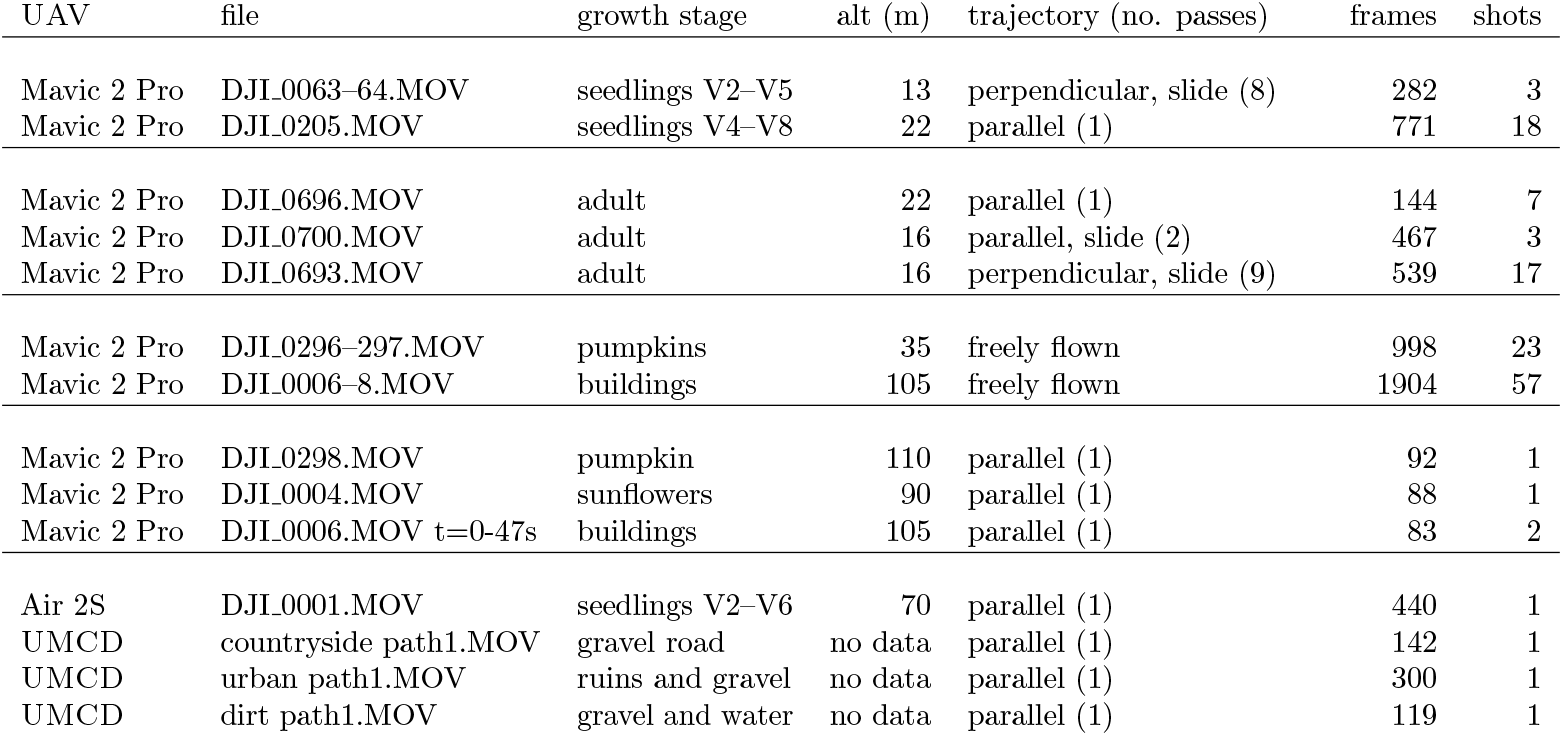
Test data used in this paper. All Mavic 2 Pro data are from the 2023 field season; the Air 2S data are from 2022. The consecutive video files arise from the UAV fragmenting the video every 3.8GB or five minutes during flight, whichever occurs first.

### 2.2 Methods

In this paper we very briefly describe our computational methods. For the details, including ablation studies, please see (Kharismawati & Kazic, 2025).

#### Dynamic sampling of video frames

We optimized frame sampling by comparing the optical flows between pairs of frames, categorizing UAV movements as translation and nontranslation. We estimated the optical flow between a base frame and a putative successor using the pixel-wise technique (Farnebäck, 2003), dividing the flows into an outer peripheral rectangle and the inner remainder, omitting the middle (see Figures 1a and 1b). The difference between the averages of the magnitudes of the outer and inner rectangles characterizes the flows. Translation is signalled by low differences in the averages while nontranslation (rotations and scaling) have higher differences. We empirically set the threshold to 5 to distinguish between translation and nontranslation.

**Figure 1:**
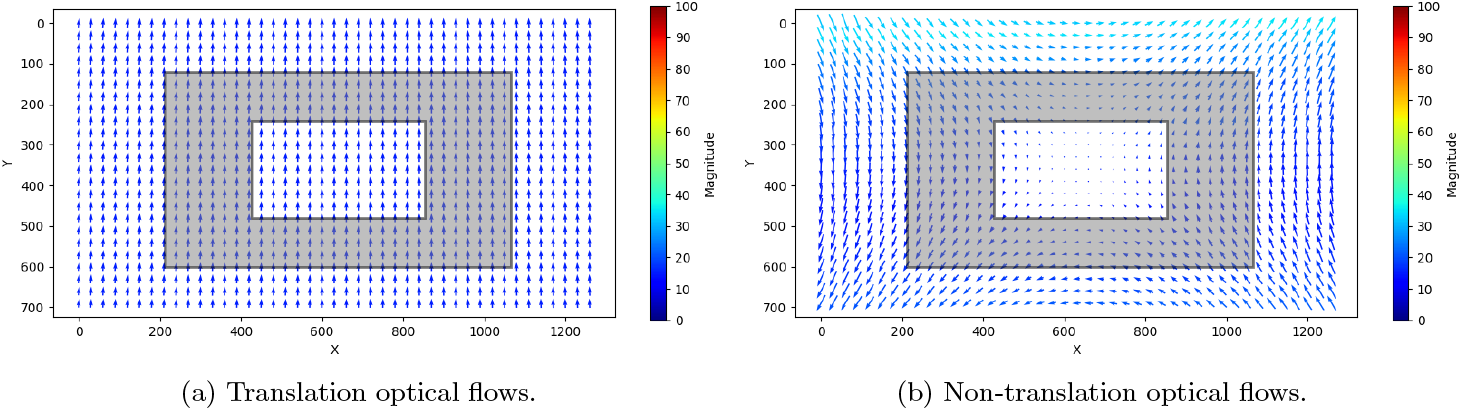
Classifying UAV movements using optical flow. Panels 1a and 1b show quiver plots of pixel-wise optical flows between two pairs of frames and the division into rectangles for classifying movements. The middle gray rectangle is not considered.

The frame sampling regimes for translations and nontranslations differ. For translations, we set the overlap threshold between a base frame and its putative successor to be between 85–98%. Overlap is measured from the displacements of the optical flows over the frame width and height, averaged over the second and third quartiles of the displacements’ distribution. If the overlap between the two frames is insufficient, the sampling rate for that region of the video is increased by half of the default rate. If the overlap is too high, the sampling rate is decreased by half until the overlap is either at least 98%, the putative successor is immediately adjacent to the base frame, or the maximum number of iterations is reached. Once the optimal successor is chosen, sampling continues using it as the new base frame. For non-translations, the default is to extract frames more frequently by increasing the sampling rate by half of the default frame sampling rate until the motion evaluates as translation.

#### Lens and gimbal calibration

Once the frames for mosaicking are chosen, we calibrate them using estimates of the camera’s extrinsic, intrinsic, and distortion parameters. These parameters were obtained for a linear, parametric model of a pinhole camera by the automatic [opencv2Camera Calibration](https://docs.opencv.org/4.x/d9/d0c/group__calib3d.html#ga69f2545a8b62a6b0fc2ee060dc30559d) subroutine using frames from orbiting the mounted checkerboard at multiple angles (Bouguet & Perona, 1998; Bouguet, 2015).

Gimbal calibration is more complex. When the UAV is in motion, it adjusts its body to maintain aerodynamic balance. For instance, during forward movement, the front propellers are positioned lower than the rear propellers. To stabilize the camera and maintain the view, the camera is mounted on a gimbal that can float in all three axes and is dynamically stabilized during flight. However, not every gimbal is perfect and wear over the course of many flights causes errors in stabilization; more modern equipment may remain more robust over the UAV’s lifetime. To address this issue for our UAV, we perform two types of corrections. The first is automatic calibration using the UAV’s software if available. The second is correction during flight, aligning the camera’s yaw and roll with the earth’s horizon. We also experimented with manual, trial-and-error post-processing calibration (see Supplemental Figure 10).

#### Homography Estimation with *CorNetv3*

We trained the new *CorNetv3* in several rounds, adding data from different field seasons, growth stages, trajectories, and movements; camera views; image calibration; and augmentation for illumination changes and noise (see Table 1). The data were balanced between adult and seedling imagery using dynamic sampling. The total training data were 117,862 pair of frames: 25,216 frames from Missouri and 33,817 frames from Indonesia, both sets paired in forward and reverse order. During training, noise and synthetic illumination changes were randomly added by altering the brightness, color, gamma, and saturation (Nguyen, Chen, Shivakumar, Taylor, & Kumar, 2018). Training on the Missouri data took approximately 116 hours on the University’s high performance computing cluster; combining the Indonesian data for continuing the training took 504 hours on our local Lambda Labs machine. Training is slowed by the low GPU memory on both machines.

#### Shot Detection

To minimize the accumulation of errors during mosaicking, we partitioned the frames into shots and mosaicked each shot separately before merging them using ASIFT (Morel & Yu, 2009). The standard graph cut algorithm works well on video that has small scene changes, such as orbits around fields, because it detects changes in perspective around a central focus (Viguier et al., 2015; Aktar et al., 2020). However, in a serpentine trajectory the perspective is relatively fixed while the focus moves. We find the changes in camera movement over some threshold by computing the change in the angle of the direction components of the optical flow vectors between each frame and its successor. A significant difference indicates a shift in the UAV trajectory, marking a potential shot boundary. For the typical serpentine shown in Panel 2b, more than 90% of the frames show angular changes of 5° or less (data not shown). The threshold is defined as the sum of the mean and standard deviation of the entire distribution of angular changes. Finally, to filter out local maxima that are too close to each other, we use `SciPy’s find_peaks` function with a specified window size. Figure 2 shows shots detected on two sample serpentine trajectories, one for adult and the other for seedling maize.

**Figure 2:**
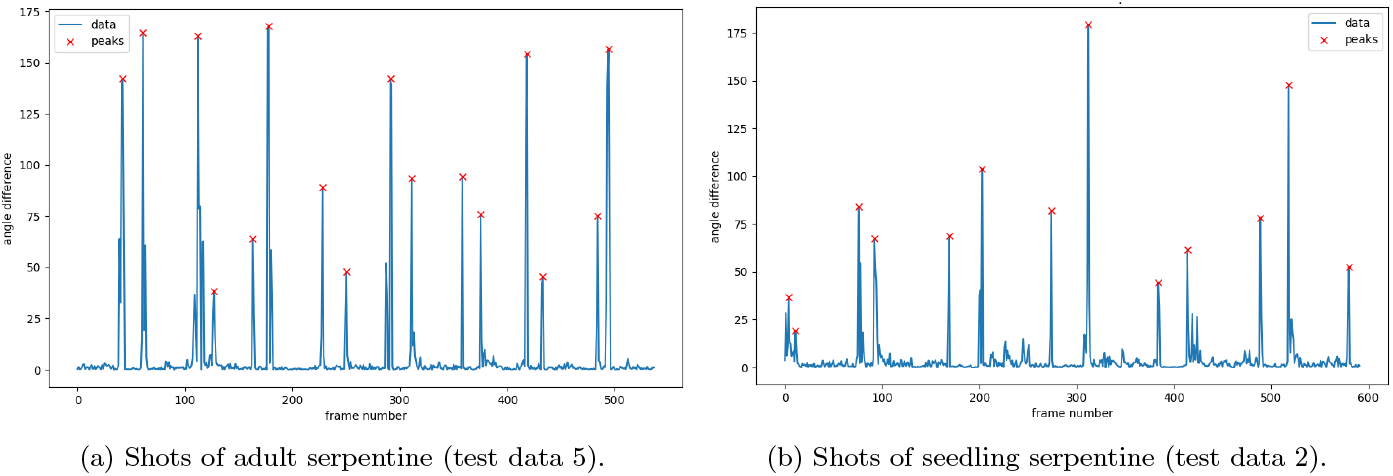
Detecting shots with optical flow. Red crosses indicate the selected shot boundaries. Panel 2a shows 16 shots detected during a serpentine with slide on adult maize, resulting in 17 mini-mosaics. Panel 2b shows 13 shots during a serpentine trajectory with slides on seedling maize, resulting in 14 mini-mosaics.

#### Mini-mosaicking

Mosaicking serpentine trajectories is more challenging than straight, few-pass trajectories because errors arise disproportionally during the turns between passes. These additional errors add to those for the subsequent pass, inflating the total accumulated error. Mosaicking *within each shot* interrupts this disproportionate error and reduces misalignments among plants. We compute the estimated homography matrix for each pair of consecutive *k* frames for that shot and accumulate in the same way as in Eqn. 1:

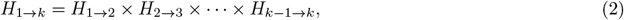

where the direction of the rightward arrow, →, denotes the forward homography estimation. Then the shot’s frames are projected onto the first, reference frame by computing the inverse matrix, *H*_1←*k*_, from *H*_1→*k*_ to warp each frame to the reference, letting the leftward arrow, ←, denote the inverted matrix. We use alpha blending where the frames overlap to replace underlying pixels with those in the final topmost frame. We use *H*_1←*k*_ to compute the projected width and height of all frames to create a canvas for the mini-mosaic. If any frame’s projected coordinates become negative, an offset matrix is computed and applied to shift everything into positive space and ensure all frames fit neatly into the canvas. Finally, each mini-mosaic is assembled into the global mosaic with ASIFT, as shown in Figure 3.

**Figure 3:**
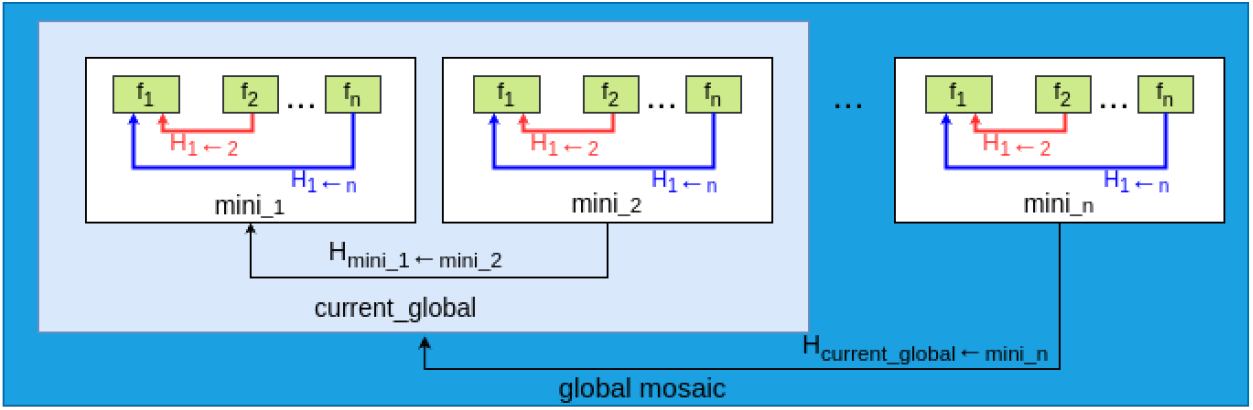
Mini-mosaicking and global mosaic assembly.

#### Evaluation

We quantitated *MaiZaic*’s accuracy several ways. First, *CorNetv3* calculates the average root mean squared error (RMSE) across all pairs of frames. Given the four corner points, *m*, of a 512 *×* 512 pixel patch in the center of the base frame, *i, CorNetv3* outputs the location of these four points projected into the putative successor frame, *i* + 1, denoted as *pred*. The ground truth of the projected points is computed with the homography matrix from ASIFT, denoted as *gt*. The average RMSE across all frames *n* was then computed as

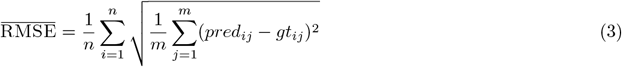

To assess the overall quality of the global mosaic, we use the geometric distortion ratio (F. Bunyak, personal communication). For a given rectangular field, this metric computes the ratio of the areas of a field’s mosaic and its ground truth bounding box. We estimated the actual area from the width and length of our fields measured by a 300’ tape measure. Since these values are very close to the measurement from Google Earth (field 27 google earth) we used the Google Earth data and derived a pixel-to-centimeter ratio (`pxtocm_ratio`) using known landmarks. When there is no landmark near a field, we used the first frame since it will be projected into the global mosaic while retaining its original geometry. The pixel area of the mosaic was estimated by clicking through the desired area with a mouse to form a polygon, then counting how many squared pixels were within the polygon using a mask. The `pxtocm_ratio` is then used to convert the mosaic’s pixel area into an actual area in square centimeters. To quantify how much the calculated mosaic area differs from the actual area, we calculated the absolute precentage error (Equation 4):

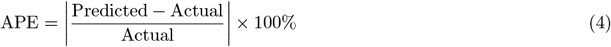

## 3 RESULTS

### 3.1 Overview

Our original *CorNet* mosaics successive pairs of frames using homography matrices estimated by a derivative of the VGG convolutional neural network trained on our maize imagery. *CorNet* can estimate homography with comparable accuracy at a tenth of the speed of ASIFT and it generalizes very well from agricultural imagery to forests, buildings, and water. However, *CorNet* was trained using uncalibrated training data from a DJI Phantom 4 of adult maize. This resulted in low accuracy when applied to the seedling datasets. Moreover, the model did not transfer well when applied to data from another drone due to different lens and gimbal errors.

To address these problems, we developed *MaiZaic*. In contrast to *CorNet*, *MaiZaic* features an end-to-end pipeline for mosaicking (Figure 4). The pipeline combines five new strategies for computing mosaics of freely-flown agricultural imagery as described in Section 2.2. Briefly, these include dynamic sampling of frames to optimize overlap; automatic calibration of lenses and gimbals; an improved deep learning network for computing homographies that is robust to growth stages, crops, trajectories, illumination changes, cameras, pilots, and vehicles (*CorNetv3*); shot detection for imagery where the focal point changes; and mini-mosaicking. Computational details will be presented in a separate paper (Kharismawati & Kazic, 2025).

**Figure 4:**
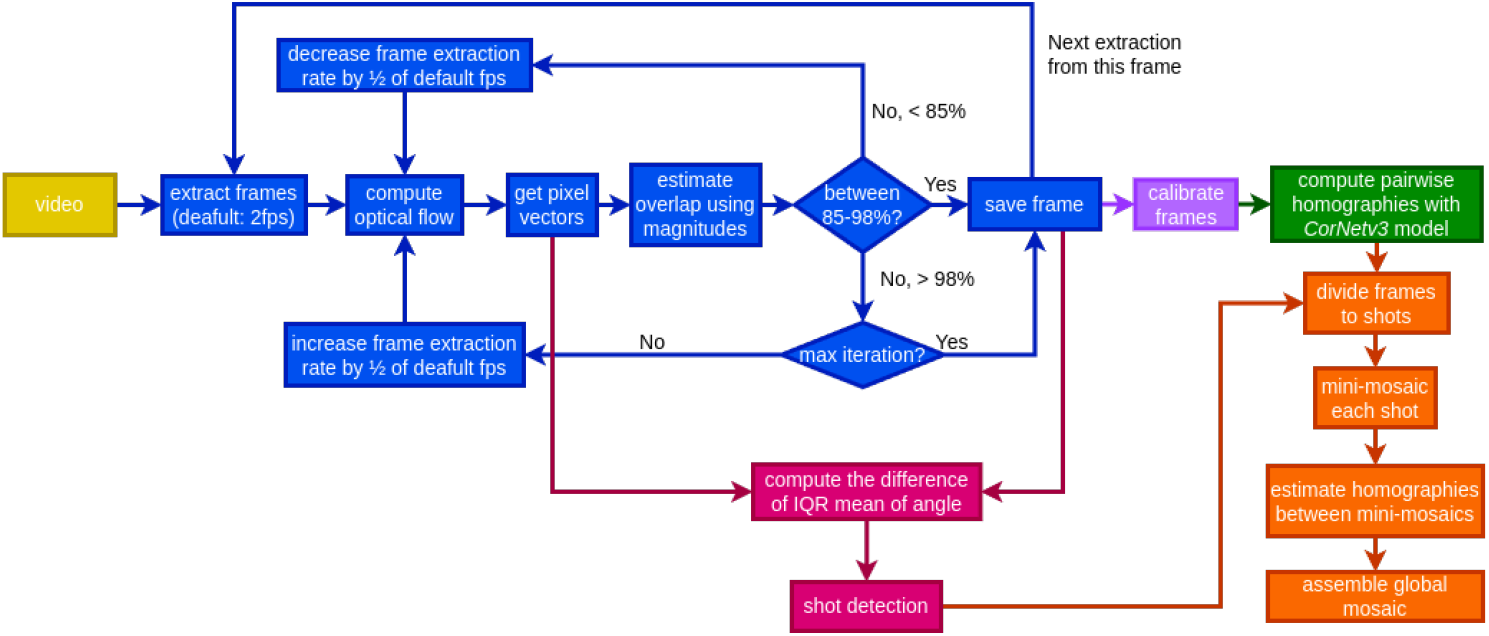
*MaiZaic* workflow. Yellow denotes the input video; blue, dynamic sampling; violet, lens and gimbal calibration; green, homography estimation with *CorNetv3*; hot pink, shot detection; and orange, mini-mosaicking, and global mosaicking.

Figure 5 compares representative mosaics computed by *MaiZaic* for seedling maize compared to the ground truth of a single frame of a rectangular field shot from a high altitude. Each mosaic was computing using the entire *MaiZaic* pipeline but using *CorNetv3*, *CorNetv2*, *CorNet*, and ASIFT to compute homographies (Morel & Yu, 2009). This strategy optimizes the input and output of each method of homography estimation for a fairer comparison. Compared to the ground truth, *MaiZaic* with *CorNetv3* shows less geometric distortion than any of the other three methods. The mosaic is essentially rectangular with gradual widening and rightward tilting as mosaicking proceeds. The *CorNetv2* mosaic is less rectangular and slants and curves more to the right. The *CorNet* mosaic is shorter, narrower at the base, and tilts to the left. Finally, the ASIFT mosaic is much longer than the ground truth and tilts more to the right than any of the other mosaics. Significantly, *MaiZaic* now mosaicks seedlings well. Similar results are seen for the less challenging case of adult maize (data not shown).

**Figure 5:**
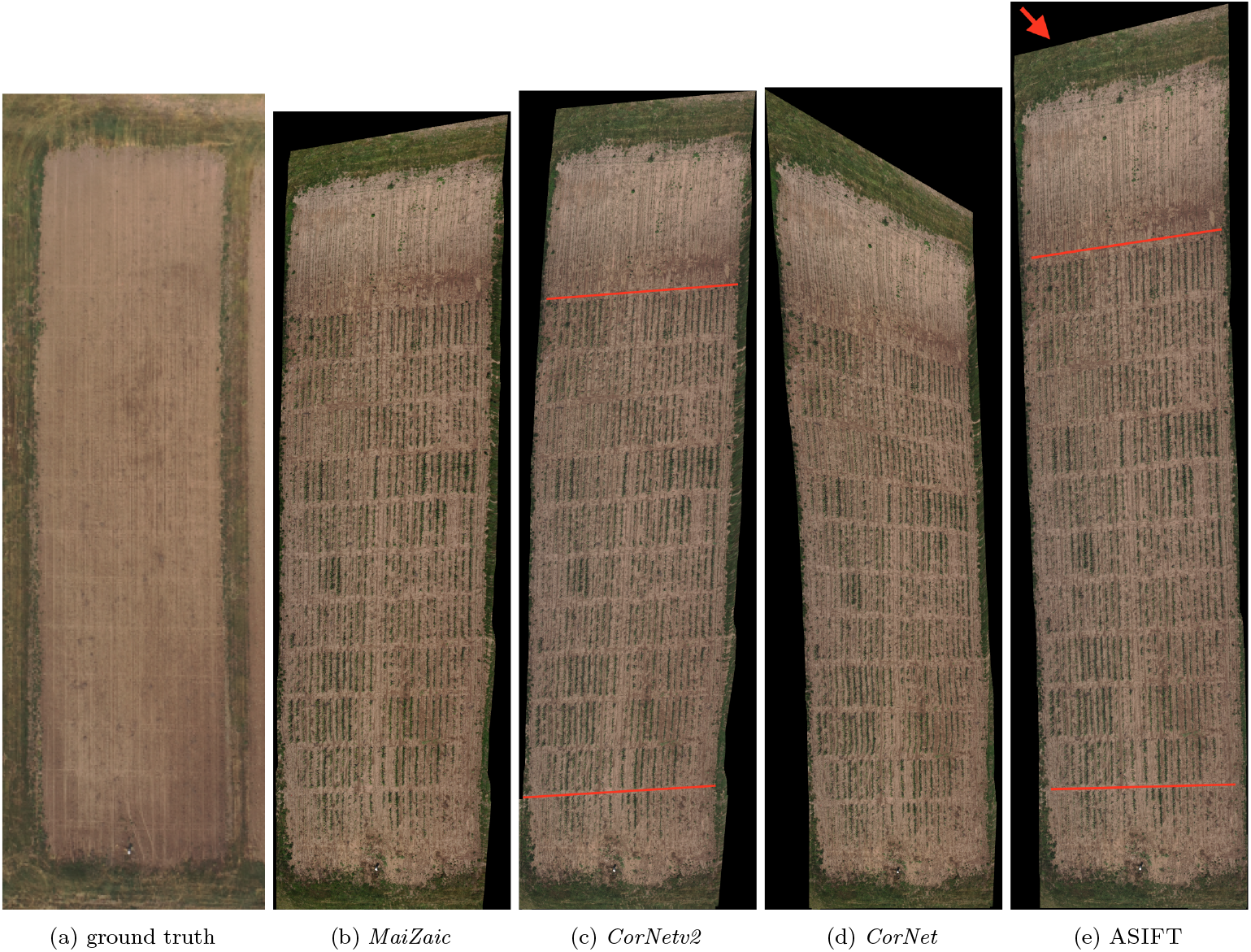
Comparison of mosaics of seedling maize computed using the entire *MaiZaic* pipeline but substituting *MaiZaic, CorNetv2*, *CorNet*, and ASIFT to compute homographies. Panel 5a is a single high-altitude image of the field before planting. The *MaiZaic* mosaic (panel 5b) was scaled so that the width of the bottom of the field matched that of the ground truth. The other mosaics are shown at the same scaling as panel 5b. The red lines mark representative alleys to aid visual comparison and the red arrow emphasizes the mosaic’s tilt. The high resolution versions of these mosaics can be found at the [MaiZaic dataset](https://drive.google.com/drive/folders/1oVcTgs3YV3AzSi2Iajl6O9g1onI7ArBN?usp=sharing).

**Figure 6:**
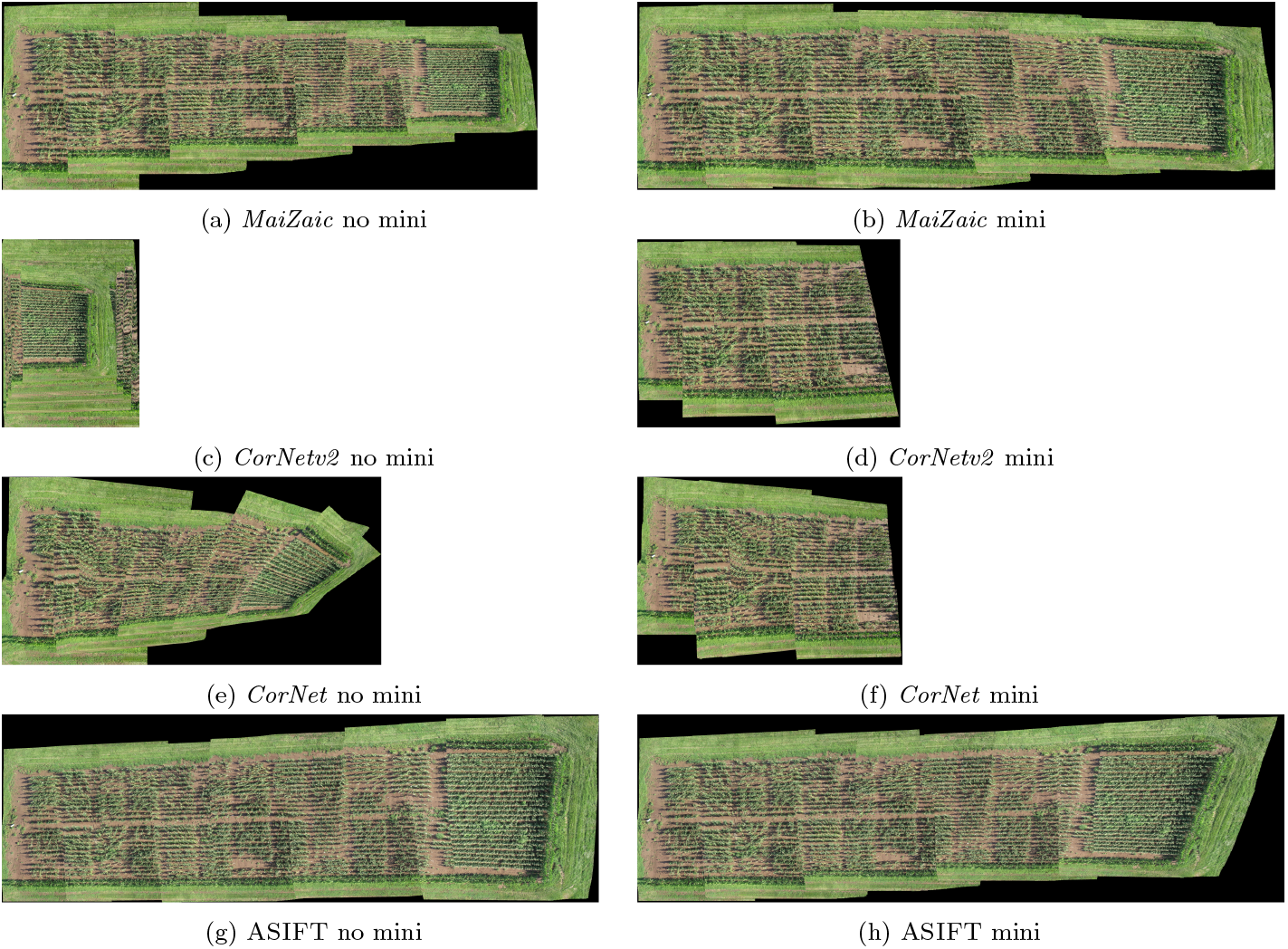
Comparison of *MaiZaic, CorNetv2*, *CorNet*, and ASIFT for mature maize with forward and backward serpentines flying perpendicular to the rows with slides and using lens calibrated frames. In the left column, panels 6a, 6c. 6e, 6g show mosaics without mini-mosaicking. In the right column, panels 6b, 6d. 6f, 6h are mosaics with shot detection and mini-mosaicking. All are the same magnification. The high resolution versions of these mosaics can be found at the [MaiZaic dataset](https://drive.google.com/drive/folders/1oVcTgs3YV3AzSi2Iajl6O9g1onI7ArBN?usp=sharing).

Table 3 compares the speed and accuracy of homography estimation for ASIFT, *CorNetv2*, and *CorNetv3*. *CorNetv3* is significantly faster than ASIFT in all trajectories and for all objects, scaling almost linearly with the number of frames in the mosaic. For all but three flights, *CorNetv3* is more accurate than *CorNetv2* for RMSE values calculated over the entire mosaic, showing relatively minor errors of 0.77 pixels for the forward trajectory and 0.85 pixels for the serpentine trajectory perpendicular to the rows with slides. *CorNetv3* also outperforms ASIFT, *CorNet*, and *CorNetv2* on mosaicking accuracy as measured by APE. For six separate flights combining these flight parameters, *MaiZaic*’s APE ranged from 1.22–5.83 compared to ASIFT’s 6.88–18.50, where lower values are better (data not shown). ASIFT’s execution time is more sensitive to the types of objects and trajectory.

**Table 3:** Execution time *per* frame and errors as a function of trajectory for ASIFT, *CorNetv2*, and *CorNetv3*, with the best results shown in bold font. *Flights* are described by object, motions relative to the field and any rows, and the number of passes; *fr* is a frame used in the mosaic; and *RMSE* is the root mean squared error in pixels for *CorNetv2* and *CorNetv3*, taking ASIFT as the ground truth (= 0). Lower values are better, as indicated by the arrows. Seedlings and adults refer to maize; the pumpkins and sunflowers were adults. All tests were run on our local Lambda Labs machine.

| *flight* | *execution time (s/fr) ↓* |  |  | RMSE ↓ |  |
| --- | --- | --- | --- | --- | --- |
|  | ASIFT | *CorNetv2* | *CorNetv3* | *CorNetv2* | *CorNetv3* |
| seedlings, linear, parallel, 1 | 13.6654 | **0.7937** | 0.8258 | **0.69** | 0.77 |
| adults, linear, parallel, 1 | 21.8918 | 0.8294 | **0.7861** | **0.77** | 0.79 |
| adults, linear, slides, parallel, 2 | 21.5515 | 0.7969 | **0.7848** | 2.04 | **0.78** |
| seedlings, linear, slides, perpendicular, 8 | 3.67482 | 0.7883 | **0.7851** | 6.63 | **0.85** |
| adults, linear, slides, perpendicular, 9 | 21.5559 | 0.7941 | **0.7835** | 6.91 | **0.87** |
| sunflowers, linear, parallel, 1 | 22.8045 | **0.8035** | 0.8165 | 0.97 | **0.72** |
| pumpkins, linear, parallel, 1 | 12.3168 | 0.7938 | **0.7920** | **0.93** | 1.02 |
| buildings, linear, parallel, 1 | 4.09687 | **0.8002** | 0.8078 | 0.97 | **0.87** |
| pumpkins, cruising | 18.0875 | 0.7884 | **0.7828** | 4.49 | **1.13** |
| buildings, cruising | 3.61733 | 0.8048 | **0.7904** | 5.62 | **1.86** |
| UMCD urban, linear, parallel, 1 | 4.45271 | **0.8141** | 0.8182 | 4.90 | **1.26** |
| UMCD dirt | 4.03142 | 0.2592 | **0.2585** | 2.89 | **1.60** |
| UMCD countryside, linear, parallel, 1 | 4.96205 | 0.8342 | **0.8331** | 4.69 | **1.28** |
| Air 2S seedling, linear, parallel, 1 | 4.32969 | 1.5538 | **1.5470** | 1.45 | **1.73** |

### 3.2 *CorNetv3* Generalizability

A perfect homography model would accurately mosaic RGB imagery from any trajectory, object type, camera, or pilot. Generalizability to trajectories enables real-time adjustment during flight. Generalizability to growth stages, crops, and objects maximizes the model’s utility. Generalizability to cameras, gimbals, rotors, and stabilization software accomodates diverse vehicles and helps compensate for misalignments that accumulate with vehicle usage. Generalizability to pilots increases operational efficiency and utilizes imagery shot by pilots with different skill levels. We assessed *CorNetv3*’s generalizability along these axes.

#### 3.2.1 Trajectories

Our freely-flown trajectories for mosaicking include forward and backward linear, sideways slide, rotation, altitude change, and free cruising components. Table 3 shows that both *CorNetv3* and *CorNetv2* execute significantly faster compared to ASIFT for all the trajectories. *CorNetv3* has the fastest execution and lowest RMSE for nearly all trajectories compared to *CorNetv2* due to the former’s more balanced training data. While *CorNetv2* performs well on simple trajectories, it struggles with complex motions because the linear trajectories make up to 75% of its training data. Our serpentine trajectory involves forward linear motion, sideways slides connecting the end of previous passes to the start of the next, and backward motion on the following passes. As illustrated in Panel 6c, *CorNetv2* misinterprets sideways motion as hovering, creating staircase-like mosaics where the new passes shrink and overlay previous ones. This reflects its tendency to memorize common trajectories so that it fails with complex movements. Retraining with balanced and varied data significantly improves accuracy (Panel 6b), enabling *CorNetv3* to achieve better RMSE and APE on complex motions while maintaining comparable execution time to *CorNetv2*.

#### 3.2.2 Objects

*CorNetv3* was trained on maize imagery, but we wanted to test its robustness for other objects that offer unique challenges, including farm buildings, sunflower fields, and pumpkin patches. Farm buildings present well-defined edges and strong features, making it easier to register their images. In contrast, sunflowers have bushier canopies but the field is continuous straight rows. The pumpkin patch is the most challenging scene, with a featureless area without structured rows and only occasional contrast from the pumpkins. *CorNetv3* shows only a slight increase in RMSE for these objects compared to maize and consistently outperforms ASIFT with respect to execution speed (Table 3). Figure 7 shows mosaics produced by *CorNetv3*. Those produced by ASIFT are qualitatively similar, but are more inaccurate in their dimensions compared to the ground truth as measured by APE (data not shown). Except for the linear trajectory over the pumpkins, *CorNetv3*’s RMSE and APE remain low and comparable to those for maize, suggesting that the model generalizes beyond maize and processes broader contextual information from the entire image patch, rather than just memorizing specific features. Since ASIFT relies on the isolated strong feature points in the frames, it can distort the mosaic if these points lie in regions of the lens, such as its periphery, that are only partially corrected by the camera parameters.

**Figure 7:**
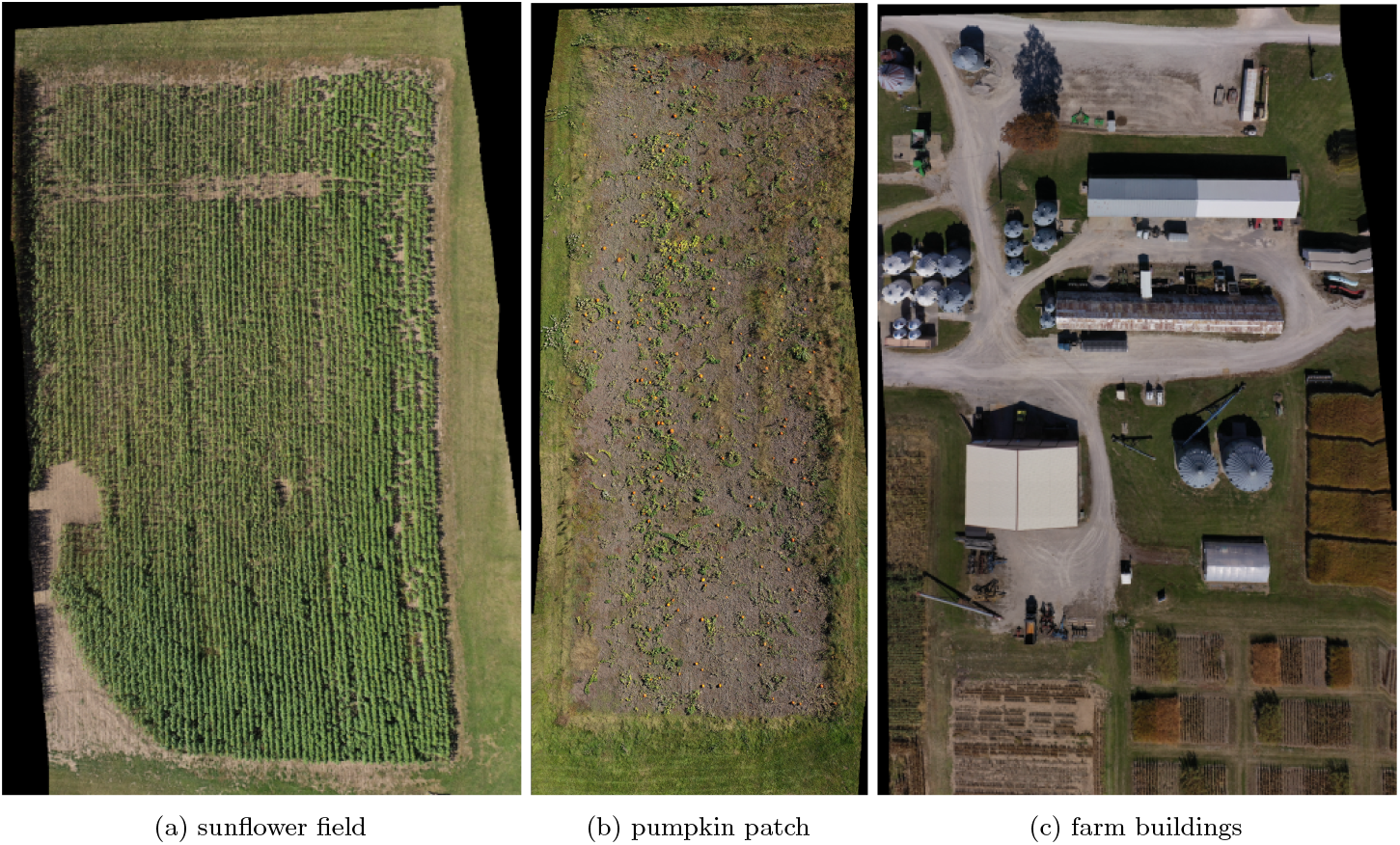
Generalizability of *MaiZaic* across diverse landscapes, underlying its versatility and broad applicability in real-world agricultural and structural settings. The high resolution versions of these mosaics can be found at the [MaiZaic dataset](https://drive.google.com/drive/folders/1oVcTgs3YV3AzSi2Iajl6O9g1onI7ArBN?usp=sharing).

#### 3.2.3 Cameras and Pilots

We evaluated *CorNetv3* ‘s generalizability across different UAV cameras and pilots. We used the UMCD dataset, which includes countryside, urban, and dirt landscapes (Avola et al., 2020). The UMCD’s camera parameters are not available for calibration, so we extracted frames at a fixed rate to mirror the experiment in (Kharismawati et al., 2020). *CorNetv3* ‘s mosaics are shown in (Figure 8) and are qualitatively comparable to those produced by ASIFT and LF-Net with excellent RMSE values (data not shown). To further assess robustness, *CorNetv3* was tested on challenging imagery from a DJI Mavic Air 2S. The flight included abrupt stops, directional changes, and altitude variations caused by a fledgling pilot. *CorNetv3* mosaicked a mature maize field better than ASIFT, though minor shrinkage toward the end of the field were observed due to gimbal error — a worsened tilt and shrinkage also seen in the ASIFT mosaics (Panel 8e). In fields with exposed soil, slight distortions appear as “wiggly” edges near the top.

**Figure 8:**
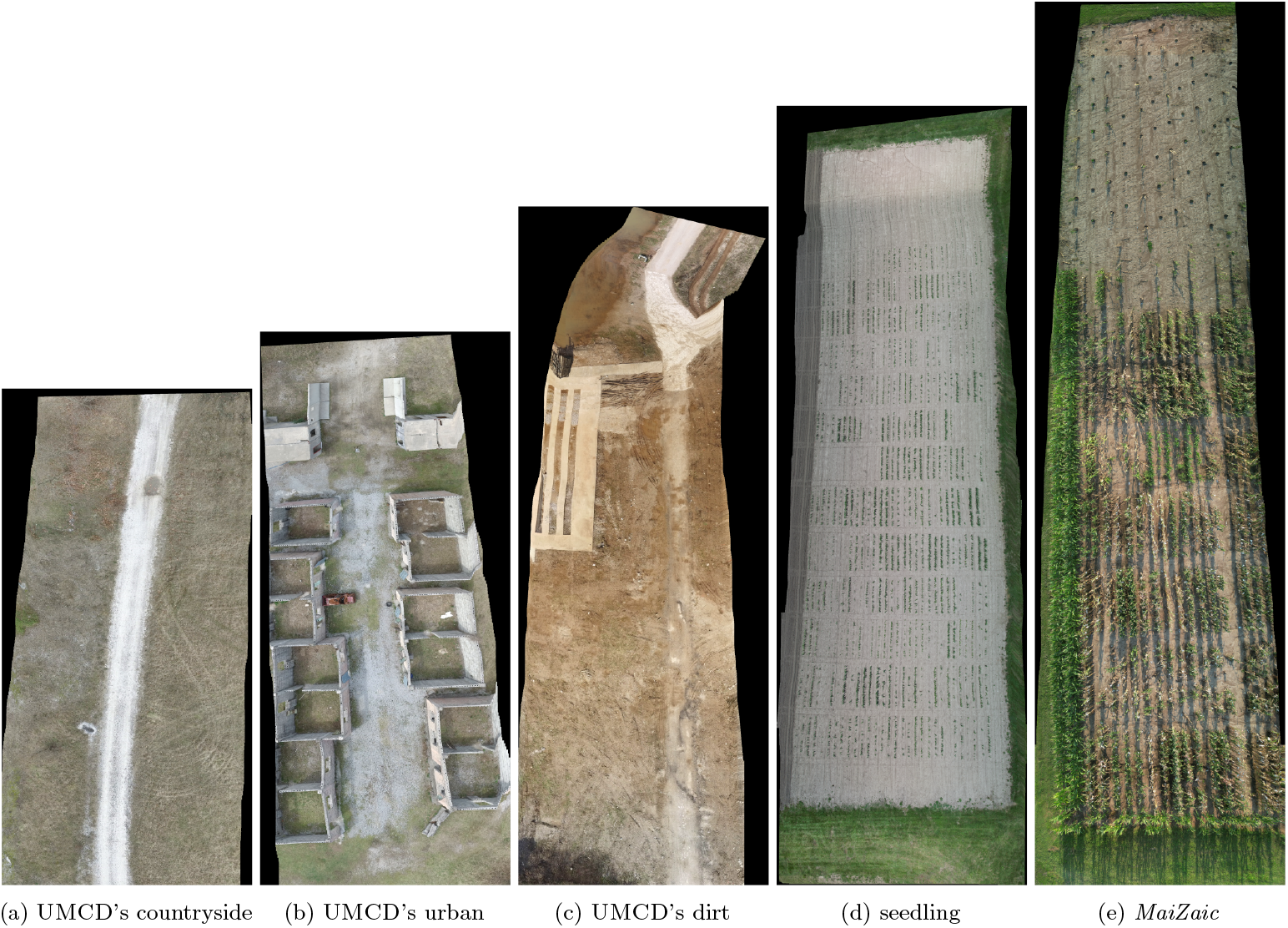
Generalizability of *MaiZaic* for different drones and pilots, showing its adaptability and practical usability for a wide range of users and UAVs. Panel 8d 8e shows seedling and adult data captured with a DJI Air 2S by a beginning pilot. The high resolution versions of these mosaics can be found at the [MaiZaic dataset](https://drive.google.com/drive/folders/1oVcTgs3YV3AzSi2Iajl6O9g1onI7ArBN?usp=sharing).

**Figure 9:**
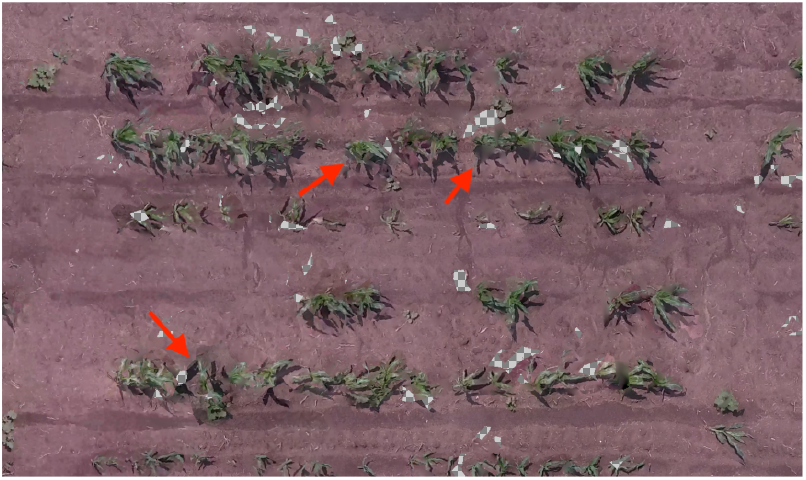
Mosaic from WebODM shows melted seedlings (a few representatives are marked with red arrows) and missing pixels (black and white check pattern).

**Figure 10:**
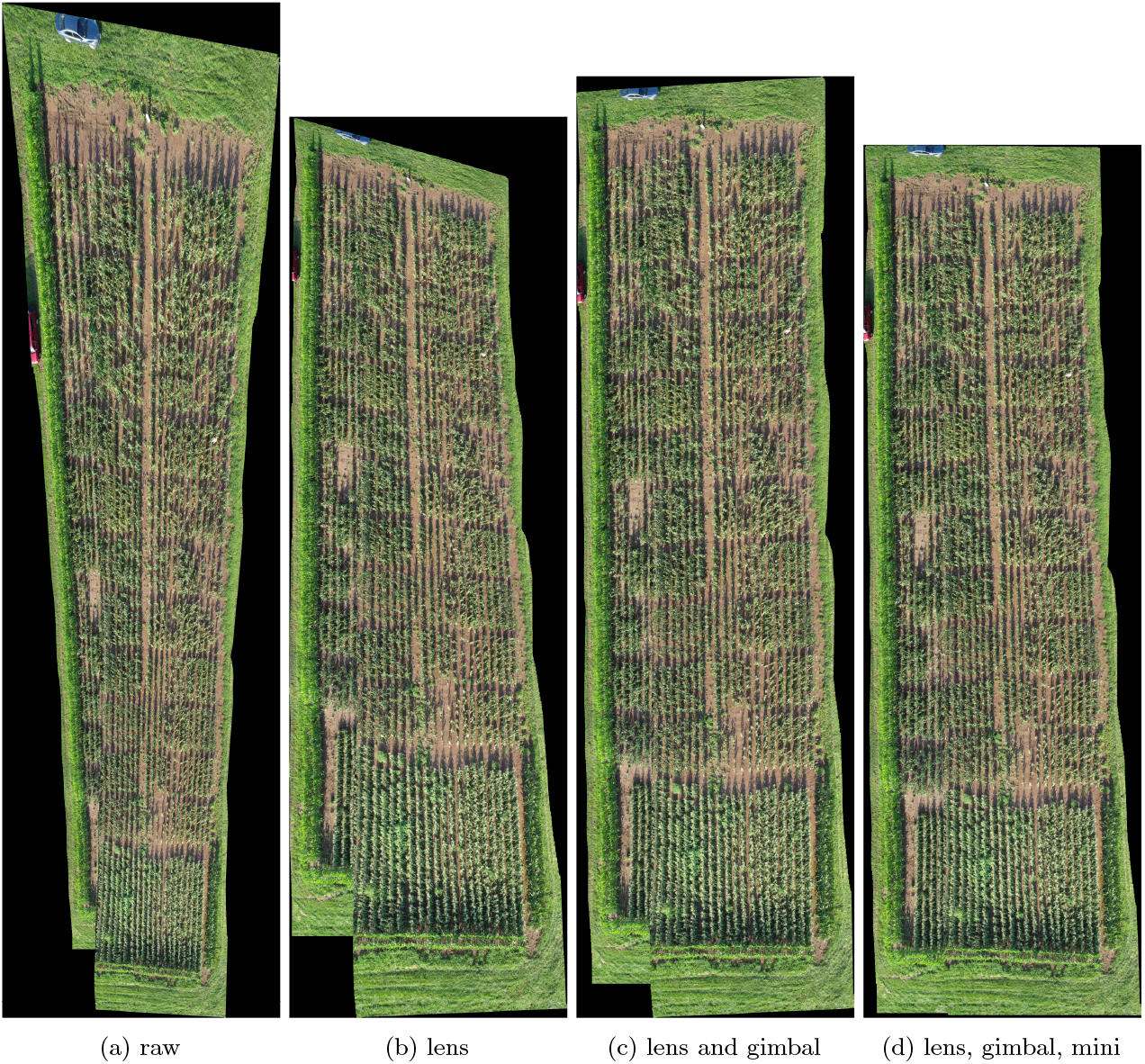
Mosaic improvement before and after lens and manual post-processing gimbal calibration and with and without mini-mosaicking with SURF.

## 4 DISCUSSION

Freely flying UAVs enables flexible, adaptive scouting for real-time crop monitoring and decision making. Without fixed trajectories, the UAVs can be immediately navigated to problematic areas, zoom in close to the plants, and capture continous high resolution imagery of smaller field sections for mosaicking to construct a view of the whole field. Mosaicking video relies on having enough overlap between adjacent frames to find similarities. This constraint motivates the common strategy of hovering over closely spaced waypoints to shoot still images. Efficient, adaptive trajectories must still satisfy this overlap constraint despite variations in altitude, speed, wind, equipment and pilots. Agricultural imagery is exceptionally challenging because distinctive features are very sparse and very similar, especially early in the season. Mature plants create repetitive patterns, complicating reliable homography estimation.

*MaiZaic* has proven to be highly effective for mosaicking these freely flown videos for several reasons. First, it extracts an optimum number of frames to maximize inter-frame overlap while avoiding excessive computational demands. Decreasing the flight speed or increasing the number of frames trivially increases the overlap and very nontrivially increases the computational time. For instance, our freely flown, unstructured videos usually capture 20 minutes at 30 fps, producing approximately 36,000 frames that require around 530GB of storage and substantial computational resources. Fixed rate sampling causes inconsistent overlap, especially during rapid maneuvers like rotations and altitude changes and causes errors in homography estimation. Our dynamic sampling approach uses Farnebäck optical flow to estimate motion between successive frames, identifying isotropic vector populations for reliable overlap. Adding more frames to low overlap regions smooths out the flows’ anisotropy and improves overlap. Thus, *MaiZaic*’s dynamic sampling ensures an overlap between 85%–98% for an optimal number of frames, reducing frame count without compromising quality.

Second, the automatic calibration capability of *MaiZaic* significantly reduces frame distortion by computing distortion matrices and intrinsic parameters for the camera lens and the UAV’s camera gimbal. Lens calibration is camera-specific and is performed using a checkerboard pattern to correct static optical distortion, while gimbal calibration addresses positional changes during flight and wear over time. Automatic gimbal calibration is performed on a flat surface using the UAV software, while in flight calibration aligns the gimbal’s roll and yaw to the horizon.

These calibrations are preferable to any post-processing corrections, which is a trial and error process that does not compensate for aerodynamic gimbal motions.

Third, the retrained *CorNetv3* is more robust to variations in growth stages, crops, flight trajectories, cameras, illumination, and pilots and so better estimates homography matrices between pairs of frames. Despite its relatively simple VGG8-like regression architecture, *CorNetv3* addresses the repetitive row patterns, homogenous plants, and significant perspective distortions caused by camera parallax. These improvements make it more resilient to challenging scenarios, including the featureless soil with repetitive patterns common in seedling data. This allows *CorNetv3* to be both faster and more accurate than ASIFT on nearly all test cases. Unlike ASIFT, which relies on the four most strongly matching points with lowest error, *CorNetv3* analyzes more complex features across the entire patch, offering greater robustness to variability in field conditions. Calibration is crucial to *CorNetv3*’s performance. Training on calibrated frames that eliminate the lens and gimbal errors prevents the model from memorizing these distortions. Furthermore, the unsupervised nature of the network allows one to incorporate more balanced and diverse data, with variations in growth stages, motions, trajectories, illumination, and camera views — all without requiring human-generated ground truth labels. The improved *CorNetv3* model strongly generalizes across different crops, flight trajectories, and UAV cameras, with the added flexibility of fine-tuning for various applications.

Fourth, dividing the frames into shots using optical flow vectors to detect changes in UAV movement addressed the challenges of error accumulation during mosaicking. By segmenting the frame sequence into smaller shots, we minimize error propagation. Unlike standard graph-cut algorithms, which struggle with constantly changing scenes, we exploit the optical flow vectors computed during dynamic sampling. Significant changes in flow were identified as shot boundaries, typically occuring when the UAV changed its direction of motion. This method was particularly effective for our serpentine flight trajectories, where flow changes happen with each pass. Occasionally, stray movements within a pass, often due to turbulence or uncontrolled updrafts and downdrafts, produced additional shots that were inconsequential for mosaicking.

Finally, mini-mosaicking these shots sharply reduces error accumulation, leading to a more accurate global mosaic for both simple and complex flight trajectories. Mosaicking aerial imagery of serpentine trajectories using homography is challenging due to matching issues: calm conditions produce excessive matches from identical plants while moving plants generate no matches. Repeated visits to the same location in serpentine trajectories further increase errors and artifactual matches. By mosaicking frames *within each shot*, we interrupt error propagation and reduce misalignment issues caused by cumulative homography matrix multiplication (Eqn. 1). However, this entails extra computation and storage for global mosaic assembly. *CorNetv3* is unsuitable for assembling the global mosaic due to its limited input patch size of 512 × 512, which leads to significant information loss when resizing mini-mosaics of varying size (*e*.*g*., 4000 × 10, 000 pixels for straight passes and 5000 × 1000 pixels for a slide). We evaluated two feature descriptors: SURF, which is faster and less accurate, and ASIFT, which is slower but more accurate. Given our focus on accuracy, ASIFT was our preferred feature descriptor. Future work will explore alternative feature descriptors such as LF-Net and SuperPoint, which may improve computational efficiency during assembly (Ono, Trulls, Fua, & Yi, 2018; DeTone, Malisiewicz, & Rabinovich, 2017).

Importantly, *MaiZaic* is robust to changes in equipment, pilots, and landscapes. The fewer the features in the landscape — such as tiny plants and the soil paths in the UMCD dirt dataset — the more any mosaicking method will struggle. *MaiZaic* demonstrates its capability to handle challenging flights, even when freely-flown by beginner pilots, smoothly mosaicking videos with sudden stops, directional shifts, speed variations, and altitude changes.

However, mosaicking free flights has some limitations that are independent of the method. Panel 8e shows the *CorNetv3* mosaic shrinks as the flight progresses and the same data processed using ASIFT exhibit a similar problem. This shrinkage increases with the age of the UAV and is independent of the training data used. Exploring alternative lens calibration patterns, such as the star and ChArUco, could yield more robust calibration parameters (An et al., 2018; Schöps, Larsson, Pollefeys, & Sattler, 2020). Additionally, *MaiZaic* requires nearly double the storage space due to its approach of partitioning each shot into subfolders to enhance computational speed, which could be optimized to eliminate this duplication. Another challenge arises during the global mosaicking assembly. As the number of mini-mosaics increases, the overlap between the new mini-mosaic and the current global mosaic diminishes, leading to potential inaccuracies in homography estimation and failed assembly. Future work could address this issue by focusing on pairwise homography estimation between mini-mosaics, particularly when dimensions are inconsistent. Despite these challenges, *MaiZaic* consistently ourperforms traditional mosaicking methods, proving to be a reliable, adaptable, and cutting-edge pipeline for UAV-based video mosaicking with broad applicability across various applications.

APE: average presentage error
ASIFT: affine scale-invariant feature transform
fps: frames *per* second
LF-NET: local feature network
RGB: red green blue color channels
RMSE: root mean squared error
SURF: speeded up robust features
UAV: unmanned aerial vehicle
UMCD: UAV mosaicking and change detection dataset
VGG8: visual geometry group (deep neural network of eight layers).

## Acknowledgments

We are deeply grateful to our Missouri maize colleagues and particularly our farm manager, Chris Browne: quality imaging depends heavily on excellent weed control, and we imaged many other fields besides our own in this work. We thank Hadi AliAkbarpour, Filiz Bunyak, Chimdi Walter Ndubuisi, Pal Palaniappan, Matthew Stanley, Dexa, and Vinny for helpful discussions.

## Data Availability Statement

All test data videos and models used in this study are publicly accessible in the MaiZaic dataset. The source code of the pipeline is available in the [Maizaic Repository](https://github.com/dek8v5/MaiZaic).

## Author Contributions

Both authors contributed to the conceptualization, methodology, validation, resources, and writing (draft and revisions). D.E.K. was responsible for software development, data collection and curation, implementation and testing, and visualization. T.K. provided supervision, project administration, and funding acquisition. Both authors have read and agreed to the published version of the manuscript.

## Supplemental Material

The supplemental material provided with this manuscript includes two figures illustrating the challenges encountered when mosaicking agricultural aerial imagery. The first is when there are visually similar plants and featureless soil; and the second is the challenges and solutions to calibrating frames that we applied during our pipeline development. Figure 9 presents a mosaic snippet generated using WebODM on seedling maize. It highlights artifacts resulting from difficulties in registering and blending similar-looking plants. The seedling maize looks melted, while the exposed soil areas contain missing pixels. Both indicate insufficient distinct features for accurate processing.

Figure 10 contextualizes the challenges associated with agricultural image mosaicking and demonstrates the iterative process of refining calibration techniques to improve results. The figure demonstrates how calibration and mini-mosaicking enhance the quality of a global mosaic constructed from serpentine trajectories. The trajectory shown is: forward parallel to row; slide right; then backward with 50% overlap between passes. In Panel 10a, raw frames were extracted using our dynamic sampling and mosaicked without any calibration or mini-mosaicking. The result shows notable distortion, especially at the top. Panel 10b illustrates the improvements achieved after lens calibration using a checkerboard pattern. While this reduces distortion significantly, some tilt remains visible at the top and misalignment issues persist at the bottom of the mosaic. Panel 10c shows the mosaic after both lens and gimbal calibration, with gimbal parameters determined by post-processing trial-and-error. This approach removes the distortion at the top but does not fully resolve the misalignment on the bottom. Finally, Panel 10d presents the mosaic generated after lens and gimbal calibration, followed by mini-mosaicking. This approach detect two shots that divide the frames into three mini-mosaics, addressing the misalignment issues and significantly improving the overall quality of the mosaic.

